# Novel genome-wide associations for suicidality in UK Biobank, genetic correlation with psychiatric disorders and polygenic association with completed suicide

**DOI:** 10.1101/451971

**Authors:** Rona J. Strawbridge, Joey Ward, Amy Ferguson, Nicholas Graham, Richard J Shaw, Breda Cullen, Robert Pearsall, Laura M. Lyall, Keira J.A. Johnston, Claire L. Niedzwiedz, Jill P. Pell, Daniel Mackay, Julie Langan Martin, Donald M. Lyall, Mark E.S. Bailey, Daniel J. Smith

**Author notes:** equal contribution (Rona J. Strawbridge, Joey Ward). Corresponding Author: Professor Daniel Smith, Address: Institute of Health and Wellbeing, University of Glasgow, Room 111, Public Health, 1 Lilybank Gardens, Glasgow, G12 8RZ, UK.

## Abstract

Background: Suicide is a major issue for global public health. ‘Suicidality’ describes a broad clinical spectrum of thoughts and behaviours, some of which are common in the general population.

Methods: UK Biobank recruited ∼0·5 million middle age individuals from the UK, of whom 157,000 completed an assessment of suicidality. Mutually exclusive groups were assessed in an ordinal genome-wide association study of suicidality: ‘no suicidality’ controls (N=83,557); ‘thoughts that life was not worth living’ (N=21,063); ‘ever contemplated self-harm’ (N=13,038); ‘an act of deliberate self-harm in the past’ (N=2,498); and ‘a previous suicide attempt’ (N=2,666). Linkage of UK Biobank to death certification records identified a small sub-group of ‘completed suicide’ (N=137).

Outcomes: We identified three novel genome-wide significant loci for suicidality (on Chromosomes 9, 11 and 13) and moderate-to-strong genetic correlations between suicidality and a range of psychiatric disorders, most notably depression (r_g_ 0·81). Higher polygenic risk scores for suicidality were associated with increased risk of completed suicide relative to controls in an independent sub-group (N=137 vs N=5,330, OR 1·23, 95%CI 1·06 to 1·41, p=0.03). Rs598046-G (chromosome 11) demonstrated a similar effect size and direction (p=0·05) within a Danish suicidality study.

Interpretation: These findings have significant implications for our understanding of genetic vulnerability to suicidal thoughts and behaviours. Future work should assess the extent to which polygenic risk scores for suicidality, in combination with non-genetic risk factors, may be useful for stratified approaches to suicide prevention at a population level.

Funding: UKRI Innovation-HDR-UK Fellowship (MR/S003061/1). MRC Mental Health Data Pathfinder Award (MC_PC_17217).

## Introduction

Suicide is a major and growing issue for global public health. Annually, approximately 800 000 people die by suicide and 20 times this number will attempt suicide during their lifetime(1). ‘Suicidality’ encompasses a broad range of experiences and behaviours, from suicidal ideas/thoughts, to acts of deliberate self-harm and suicide attempts, occurring along a spectrum towards completed suicide(2). Some components of suicidal thoughts and behaviours, such as feeling that life is not worth living or contemplating self-harm, are relatively common in the general population (as well as in clinical populations). Suicidality can therefore be considered a complex trait that fits within a Research Domain Criteria (RDoC) approach because it cuts across traditional psychiatric diagnostic classifications.

Pathways to completed suicide are complex and multifactorial(3). Suicidal thoughts and actions are a consequence of a dynamic interplay between genetic, other biological, psychiatric, psychological, and a wide range of important social, economic and cultural factors(4). Clinically, deliberate self-harm (DSH) is a major risk factor for subsequent suicidal behaviour. It is also recognised that substance abuse-related disorders and mood disorders are particularly associated with suicide risk(5). Similarly, early adversities such as childhood sexual abuse(6), maladaptive parenting(7) and parental loss(8) all contribute to suicidal thoughts and behaviours, either directly or by increasing the risk of psychiatric disorders(9, 10). Personality-level traits such as neuroticism, impaired decision-making and sensitivity to negative social stimuli(11) also contribute to suicidality.

Family, adoption and twin studies suggest a heritability estimate for suicidal behaviour of approximately 45%(12), thus suicidality is amenable to genetic investigation. Heritability estimates for less clearly defined phenotypes (such as suicidal thoughts) are difficult to establish(13). It is clear, however, that genetic predisposition plays a role alongside individual and social factors. Genome-wide association studies (GWAS) may offer some insight into the biological basis of suicidality but findings to date have been limited: A recent GWAS of suicide attempts in ∼50 000 individuals with and without psychiatric disorders identified a number of suggestive loci(14) and a single GWAS significant finding (on Chromosome 2)(15) was identified in early studies, but this has not been replicated, perhaps due to subsequent under-powered studies and heterogeneous and highly selected clinical samples.

Our goals in this study were: to identify genetic variants associated with broadly-defined suicidality in 122 935 participants of the UK Biobank cohort; to assess for genetic correlations between suicidality and a range of psychiatric disorders; and to determine whether increased genetic burden for suicidality was associated with both psychiatric disorders and completed suicide. In secondary analyses, mindful that not all DSH behaviours carry active suicidal intent, we also conducted additional separate GWAS analyses of DSH and suicidal ideation/attempts (SIA).

## Materials and methods

### Sample description

UK Biobank is a large general population cohort. Between 2006 and 2010, approximately 502,000 participants (age range 37-73 years) were recruited from around the UK (excluding Northern Ireland) and attended one of 22 assessment centres across the UK(16, 17). Comprehensive baseline assessments included sociodemographic characteristics, cognitive abilities, lifestyle and measures of mental and physical health status (Supplementary Methods and Supplementary Figure 1). Only white British participants were included in the current analysis. Informed consent was obtained by UK Biobank from all participants. This study was carried out under the generic approval from the NHS National Research Ethics Service (approval letter dated 13 May 2016, Ref 16/NW/0274) and under UK Biobank approval for application #6553 “Genome-wide association studies of mental health” (PI Daniel Smith).

### Suicidality phenotypes

Suicidality categories were based on four questions from the self-harm behaviours section of the online mental health (*‘Thoughts and Feelings’)* questionnaire: (http://biobank.ctsu.ox.ac.uk/crystal/label.cgi?id=136 and Supplementary Methods(18)). Linkage to death certification (until February 2016) identified a sub-group of participants classified as ‘completed suicide’ (N=137). Non-overlapping categories of increasing severity of suicidality were derived: ‘no suicidality’ controls; ‘thoughts that life was not worth living’; ‘ever contemplated self-harm or suicide’; ‘acts of deliberate self-harm not including attempted suicide’; ‘attempted suicide’; and ‘completed suicide’. Note that ‘completed suicide’ was not used in the ordinal GWAS but rather was used as a separate sub-group to test for association with genetic loading for suicidality. Participants were classified based on the most extreme form of suicidality that they reported and within the ‘no suicidality’ group if they responded negatively to all self-harm and suicidality questions. Those who preferred not to answer any of the questions were excluded from analysis (0·7%). Completed suicide was defined as primary cause of death by intentional self-harm (ICD codes X60-X84).

### Genotyping, imputation and quality control

In July 2017 UK Biobank released genetic data for 487 409 individuals, genotyped using the Affymetrix UK BiLEVE Axiom or the Affymetrix UK Biobank Axiom arrays (Santa Clara, CA, USA)(17). These arrays have over 95% content in common. Pre-imputation quality control, imputation and post-imputation cleaning were conducted centrally by UK Biobank (described in the UK Biobank release documentation(16, 17). Fully imputed genetic data released in March 2018 were used for this study.

### Ordinal GWAS of suicidality, DSH and SIA

For each GWAS, we excluded at random one person from each related pair of individuals with a kinship coefficient > 0·042 (second cousins) that have valid phenotypes, therefore the number of controls is different for each analysis (Supplementary Methods and Supplementary Figures 1 and 2).

The primary GWAS included 122,935 individuals. Of these, 83,557 were classified as controls (category 0), 21,063 were classified into ‘thoughts that life is not worth living’ group (category 1), 13,038 into the ‘thoughts of self-harm’ group (category 2), 2,498 into the ‘actual self-harm’ group (category 3), and 2,666 into the ‘attempted suicide’ group (category 4).

For the secondary analyses, two further GWAS were conducted. For DSH, the categories of controls, (N=84,499), “thoughts of self-harm’ (N=13,203) and ‘actual self-harm’ (N=2,532) were assessed. For SIA, the categories of controls (N=84,167), ‘thoughts that life is not worth living’ group (N=21,234) and ‘attempted suicide’ (N=2,689) were assessed.

Analyses were performed in R (Version 3.1) using the clm function of the ordinal package (19) treating the multilevel suicidality, DSH or SIA outcome variable as an ordinal variable. Models were adjusted for age, sex, genotyping chip and UK Biobank-derived genetic principal components (GPCs) 1-8. For sensitivity analyses, additional adjustment for psychiatric diagnosis (defined as self-reported BD, GAD and MDD) was also applied. Genome-wide significance was set at *P*<5×10^-8^ and plots were generated using FUMA(20).

### Polygenic Risk Score (PRS) variables for suicidality, mood disorders and related traits

PRSs were calculated from the primary ordinal suicidality GWAS summary statistics. SNPs were included in the PRS if they met *P*-value thresholds of *P*<5×10^-8^, *P*<5×10^-5^, *P*<0·01 *P*<0·05, *P*<0·1 or *P*≤0·5 (Supplementary Methods). PRS deciles were computed using STATA (version 12, STATACorp) and modelling of associations between the PRS and completed suicide was analysed with logistic regression, adjusting for age, sex, chip and GPCs 1-8. In this analysis, cases were individuals classified as ‘completed suicide’ (n=137), and controls were those recorded as category 0 in the ordinal variable but who had been excluded from the GWAS due to relatedness (n=5,330). Associations between the PRS and risk of mood disorders and related traits were also assessed (Supplementary Methods). The traits tested (BD, MDD, mood instability, and risk-taking behaviour) were selected based on prior evidence (epidemiological or clinical) of relevance to suicidality, therefore the threshold for significance was set at *P*<0·05.

### SNP heritability and genetic correlation analyses

Linkage Disequilibrium Score Regression (LDSR)(21) was used to estimate the SNP heritability (h^2^SNP) of ordinal suicidality, DSH and SIA. LDSR was also used to calculate genetic correlations with suicide attempt, psychiatric disorders and related traits (Supplementary Methods). The resulting genetic correlation *P*-values were false discovery rate (FDR)-corrected to compensate for multiple testing.

### Gene-based analysis

The ordinal GWAS results were also considered under a gene-based approach, using MAGMA(22), as implemented in FUMA(20).

### Exploration of known biology

The Variant Effect Predictor web-based tool(23), GTEx database(24) and BRAINEAC dataset (http://braineac.org) were interrogated to try to identify genes (based on expression quantitative trait loci, eQTLs) or mechanisms through which associated SNPs might be acting (Supplementary Methods). The GWAS catalogue (https://www.ebi.ac.uk/gwas) and NCBI Gene https://www.ncbi.nlm.nih.gov/gene) were queried for suicidality-associated SNPs and genes.

## Results

### Sociodemographic characteristics

Sociodemographic, clinical and health behaviour measures for each of the suicidality categories are shown in Supplementary Table 1. As expected, a gradient of increasing suicidality was found for increasing levels of social deprivation, living alone, current or previous smoking, parental depression and chronic pain. There were also substantial differences by sex: females accounted for 68·3% of those who reported attempted suicide but only 27·6% of completed suicides. A large proportion of those who had attempted suicide (85·1%) had a history of MDD, compared to only 14·9% of controls. Similarly, 75·8% of those with a suicide attempt had reported childhood trauma, compared to 39·0% of controls.

### Primary ordinal GWAS of suicidality

The results of the ordinal GWAS of suicidality are presented in Table 1, Supplementary Table 2 and Figure1A. The GWAS results showed some inflation of the test statistics from the null (λ_GC_ =1·16, Figure 1A, inset) but this was not significant given the sample size used (λ_GC_ 1000=1·004). LDSR demonstrates that polygenic architecture, rather than unconstrained population structure, is the likely reason for this (LDSR intercept = 1·02, SE=0·0075). SNP heritability was estimated by LDSR as being 7·6% (observed scale h^2^SNP = 0·076; SE=0·006).

**Figure 1:**
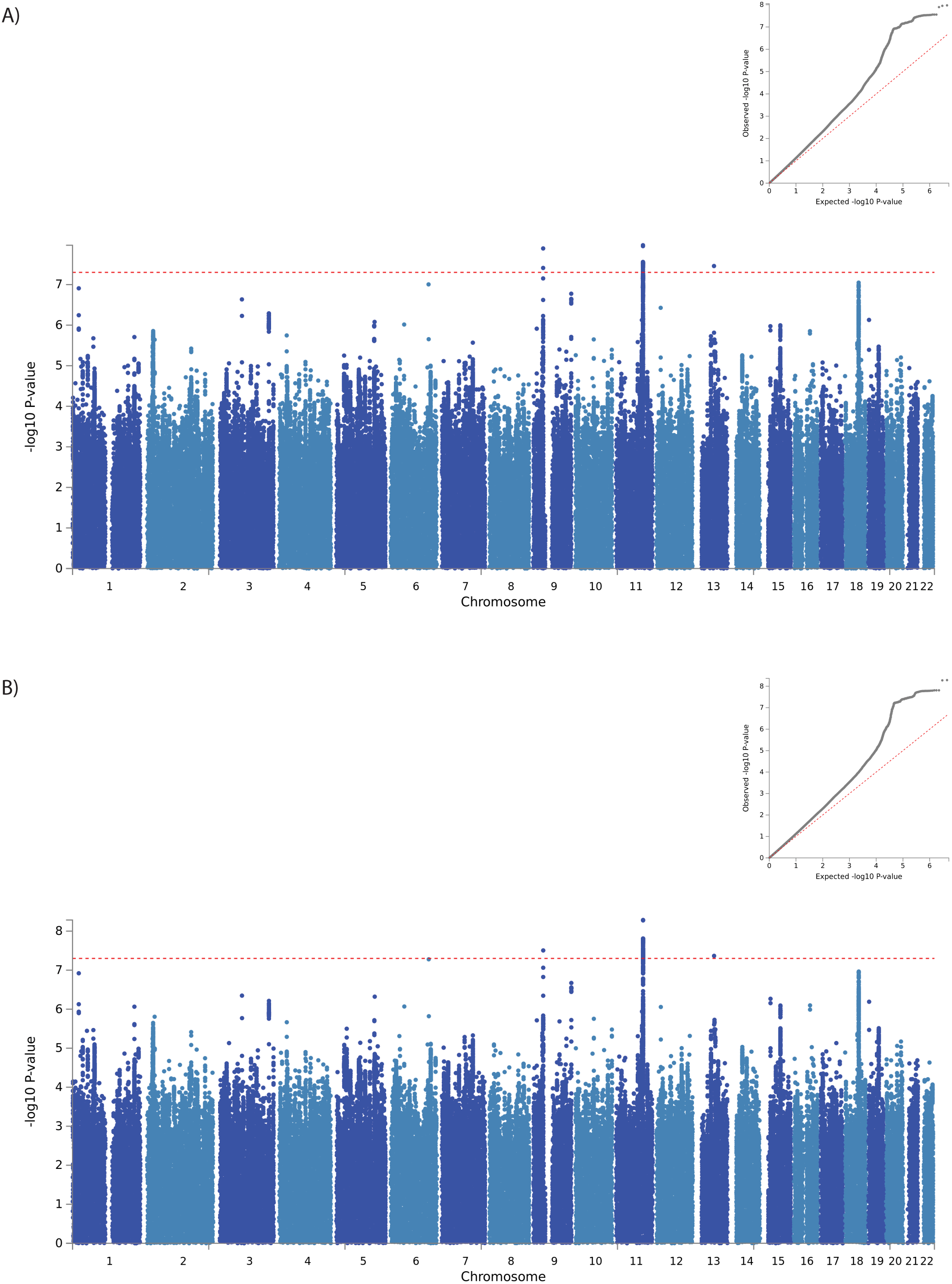
Manhattan plot of GWAS of ordinal suicidality in UK Biobank (N=122 935): A) adjusted for age, sex, genotyping chip and population structure, B) adjusted for age, sex, genotyping chip, population structure and psychiatric disorders. Dashed red line = genome wide significance threshold. Inset: QQ plot for genome-wide association with DSH. Red line = theoretical distribution under the null hypothesis of no association

**Table 1:** **Lead SNPs at loci associated with ordinal suicidality at GWAS significance**

| Analysis | SNP | CHR | POS | A1 | A2 | BETA | SE | *P* | A1F<sup>a</sup> | SNPs_gwas<sup>b</sup> | SNPs_sugg<sup>b</sup> |
| --- | --- | --- | --- | --- | --- | --- | --- | --- | --- | --- | --- |
| Ordinal Suicidality | rs62535711 | 9 | 37174829 | T | C | 0.105 | 0.018 | 1.29E-08 | 0.056 | 15 | 57 |
|  | rs598046 | 11 | 99516468 | T | G | 0.053 | 0.009 | 1.07E-08 | 0.319 | 34 | 370 |
|  | rs7989250 | 13 | 64900801 | A | C | -0.052 | 0.009 | 3.49E-08 | 0.322 | 1 | 5 |

Where: A1, effect allele; A2, other allele; A1F, effect allele frequency; <sup>a</sup> calculated in whole cohort; <sup>b</sup> within the region defined by suggestive significance ( $P < 1 \times 10^{-5}$ ); Aligned to Human Genome assembly GRCh37, Chr9 locus, 9:36999369-37360767; Chr11 locus, 11:99392678-99588751; Chr13 locus, 13:64900801-65036; Chr6 locu, 6:140442326-140895470; SIA, suicidal ideation or attempt.

We identified three independent loci associated with suicidality (Table 1, Supplementary Table 2 and Figure 1A): one on chromosome 9 (index SNP rs62535711, Figure 2A) within the gene ZCCHC7; a second on chromosome 11 (index SNP rs598046, Figure 2B) located within *CNTN5;* and a third on Chr.13 (index SNP rs7989250, Figure 2C). Conditional analyses (Supplementary Methods) in which the lead SNP was included as a covariate demonstrated no significant secondary association signals at these loci (the most significant SNP on Chr.9 was rs999510, *P*=0·0008; that on Chr.11 was rs608820, *P*=0·0005; and that on Chr.13 was rs9564176, *P*=0·003). Adjustment of the GWAS for psychiatric disorders had little effect on the observed associations (Figure 1B and Supplementary Table 2). Effect allele frequencies by suicidality category are presented in Supplementary Table3.

**Figure 2:**
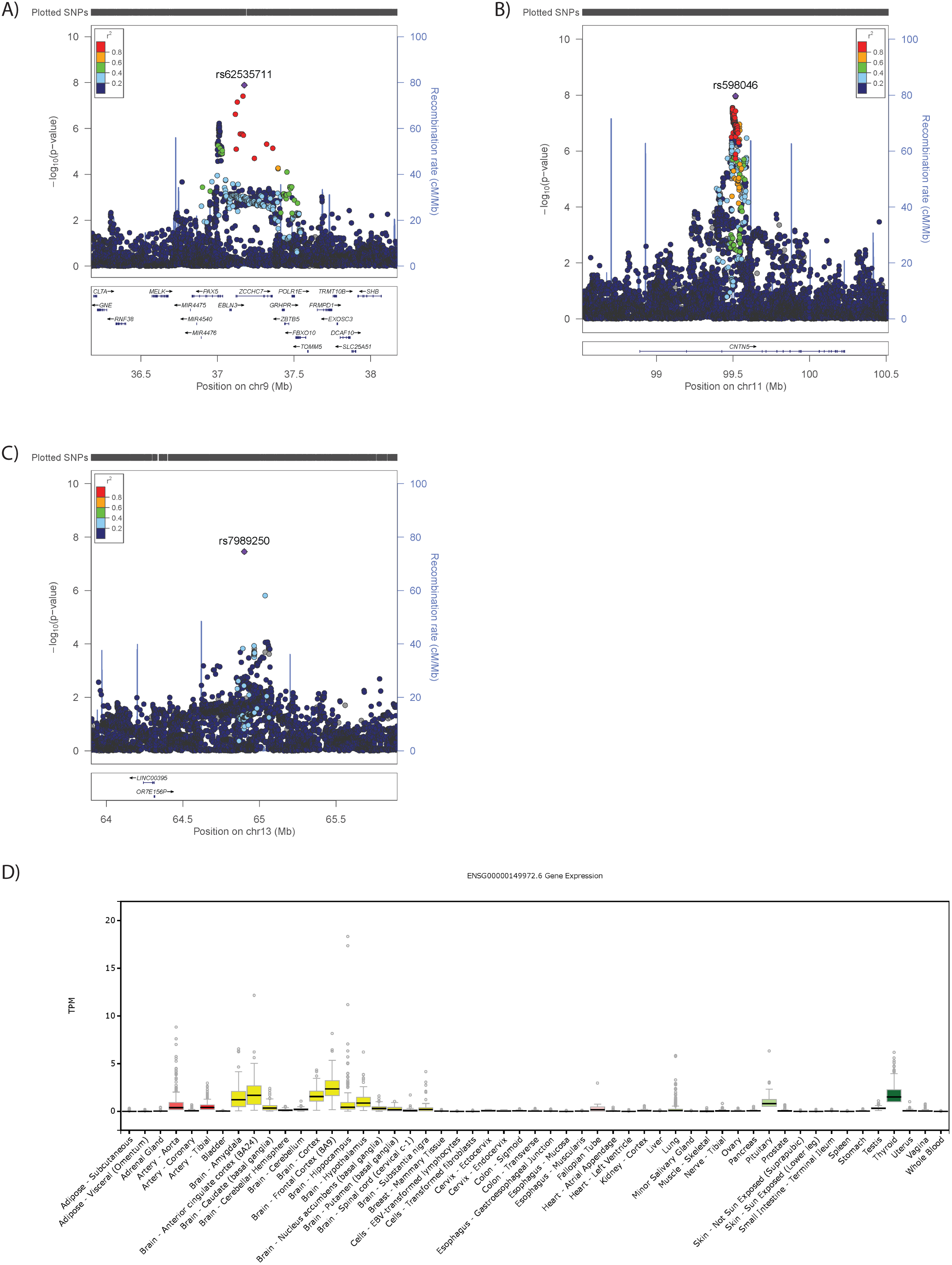
Regional plots for GWAS significant loci: A) *ZCCHC7* locus on Chr9, B) *CNTN5* locus on Chr11, C) Chr13 locus, where: SNPs (each point) are aligned according to position (X axis) and strength of association (Y Axis, left); Purple colouring indicates the index SNP, with other colours representinglinkage disequilibrium (r^2^) with the index SNP, as per the colour key; Rates of DNA recombination are presented as a pale blue line graph in the background (Y axis, right); Genes are presented by location (X axis) and direction of transcription (arrows). D) Tissue expression profile of *CNTN5*, where tissues are arranged alphabetically along the X axis and expression level is provided on the Y axis. Box plots represent median and interquartile range, with error bars demonstrating 1.5x the interquartile range and dots representing outliers.

Within the recently-reported GWAS of suicide attempt(14), rs62535711 and rs7989250 were non-significant (p=0·278 and p=0·152 respectively) whilst rs598046 was borderline nominally significant (G allele, Beta 0·041, Se 0·021 p=0·051).

### Association between genetic loading for suicidality and ‘completed suicide’

Demographic characteristics of the controls in this analysis were comparable to those included in the GWAS (Supplementary Table 1). Individuals within the completed suicide group followed the pattern of increasing deprivation, more childhood trauma and higher prevalence of mood disorders observed across the categories of increasing suicidality (Supplementary Table 1). We investigated whether greater genetic loading for suicidality, indexed by PRS for suicidality, was associated with completed suicide. Overall, higher values of suicidality PRS were associated with an increased risk of completed suicide at all but one of the PRS significance thresholds assessed (for example, *P-*threshold 0·05: OR 1.23, 95%CI 1·06-1·41, FDR p=0·04, Table 2).

**Table 2:** **Increasing burden of suicidal behaviour-associated variants significantly associated with completed suicide**

| Threshold | OR | L95 | U95 | p | FDR-adj p |
| --- | --- | --- | --- | --- | --- |
| $5 \times 10^{-8}$ | 1.07 | -0.10 | 0.24 | 0.410 | 0.410 |
| $5 \times 10^{-5}$ | 1.20 | 0.01 | 0.35 | **0.041** | **0.049** |
| 0.01 | 1.22 | 0.02 | 0.38 | **0.026** | **0.390** |
| 0.05 | 1.26 | 0.05 | 0.41 | **0.011** | **0.034** |
| 0.1 | 1.27 | 0.06 | 0.42 | **0.008** | **0.034** |
| 0.5 | 1.25 | 0.03 | 0.41 | **0.021** | **0.039** |

Where: Threshold, GWAS $P$ threshold of the SNPs included in the suicidality PRS; OR, odds ratio; L95, lower 95% confidence interval; U95, upper 95% confidence interval; Z, test statistic, p, p value for analysis; FDR-adj p, false discovery rate adjusted p. Significance was set at $P < 0.05$

### Genetic loading for suicidality and mood disorders

PRS for suicidality demonstrated consistent significant associations with mood disorders (BD and MDD) and related traits (mood instability, neuroticism, and risk-taking propensity), across most of the significance thresholds assessed (Supplemental Table 4).

### Secondary GWAS analyses

The GWAS of DSH (controls, contemplates self-harm and actual self-harm) identified no SNPs reaching GWAS significance (Supplementary Figure 3A). Adjustment for psychiatric diagnosis had little impact on these results (Supplementary Figure 3B): lead SNP rs4521702-T, Beta −0·01162, Se 0·0218, *P*=9·56×10^-8^; and Beta −0·01159, Se 0·0218, *P*=1·10×10^-7^, without and with adjustment for psychiatric diagnosis, respectively.

The GWAS of SIA (controls, suicidal ideation, suicide attempts) identified no SNPs at GWAS significance (Supplementary Figure 4A), however adjustment for psychiatric disorders identified a singleton SNP at GWAS significance (Supplementary Figure 4B-C and Supplementary Table 2). In the recent GWAS of suicide attempt (14), this SNP, rs116955121, was non-significant (p=0·509).

### Genetic correlation analyses

When considering the whole genome (rather than SNPs selected for association with suicidality, as is the case for the PRS), we observed significant genetic correlations between suicidality (primary analysis) and attempted suicide as well as all of the major psychiatric disorders and traits assessed (Table 3). The strongest genetic correlations were observed for MDD (r_g_ 0·81), neuroticism (r_g_ 0·63) and mood instability (r_g_ 0·50). DSH demonstrated similar genetic correlations with attempted suicide psychiatric disorders and related traits as those observed for suicidality (Table 3). In contrast, for SIA, significant genetic correlations were observed only for MDD, schizophrenia, neuroticism and mood instability (Table 3).

**Table 3:** **Genetic correlations of suicidality with psychiatric disorders and related traits**

| Trait | Suicidality (-MH) |  |  |  |  | DSH (-MH) |  |  |  |  | SIA (-MH) |  |  |  |  |
| --- | --- | --- | --- | --- | --- | --- | --- | --- | --- | --- | --- | --- | --- | --- | --- |
| | $r_g$ | se | z | p | FDR-P | $r_g$ | se | z | p | FDR-P | $r_g$ | se | z | p | FDR-P |
| Attempted suicide | 0.57 | 0.096 | 5.98 | **2.23E-09** | **6.24E-09** | 0.624 | 0.106 | 5.8897 | **3.87E-09** | **9.03E-09** | -0.012 | 0.182 | -0.068 | 9.46E-01 | 9.83E-01 |
| MDD | 0.81 | 0.04 | 18.66 | **1.01E-77** | **1.41E-76** | 0.79 | 0.05 | 14.48 | **1.68E-47** | **2.35E-46** | 0.46 | 0.11 | 4.07 | **4.76E-05** | **2.22E-04** |
| Neuroticism | 0.63 | 0.04 | 16.48 | **5.51E-61** | **3.86E-60** | 0.57 | 0.05 | 12.09 | **1.26E-33** | **8.82E-33** | 0.56 | 0.12 | 4.77 | **1.80E-06** | **1.26E-05** |
| Mood Instability | 0.50 | 0.03 | 16.06 | **4.53E-58** | **2.11E-57** | 0.43 | 0.04 | 11.73 | **8.50E-32** | **3.97E-31** | 0.53 | 0.11 | 5.01 | **5.42E-07** | **7.59E-06** |
| Schizophrenia | 0.32 | 0.04 | 8.70 | **3.19E-18** | **1.12E-17** | 0.31 | 0.04 | 7.02 | **2.25E-12** | **7.88E-12** | 0.28 | 0.09 | 2.96 | **3.06E-03** | **1.07E-02** |
| Bipolar disorder | 0.27 | 0.05 | 5.28 | **1.26E-07** | **2.94E-07** | 0.27 | 0.06 | 4.34 | **1.45E-05** | **2.26E-05** | 0.19 | 0.12 | 1.55 | 1.21E-01 | 3.39E-01 |
| Risk-taking behaviour | 0.20 | 0.04 | 5.05 | **4.44E-07** | **7.77E-07** | 0.25 | 0.04 | 6.00 | **1.93E-09** | **5.40E-09** | 0.00 | 0.08 | 0.02 | 9.83E-01 | 9.83E-01 |
| Anxiety disorder | 0.75 | 0.17 | 4.35 | **1.38E-05** | **2.15E-05** | 0.87 | 0.20 | 4.41 | **1.03E-05** | **1.91E-05** | 0.31 | 0.30 | 1.03 | 3.05E-01 | 5.34E-01 |
| ADHD | 0.21 | 0.05 | 4.29 | **1.79E-05** | **2.51E-05** | 0.23 | 0.05 | 4.40 | **1.09E-05** | **1.91E-05** | -0.12 | 0.11 | -1.17 | 2.44E-01 | 5.20E-01 |
| PTSD | 0.42 | 0.16 | 2.70 | **6.97E-03** | **8.87E-03** | 0.52 | 0.18 | 2.82 | **4.82E-03** | **6.75E-03** | 0.16 | 0.28 | 0.57 | 5.70E-01 | 7.98E-01 |

Where: $r_g$ , genetic correlation; se, standard error of genetic correlation; z, test statistic, P, P value for analysis; FDR-P, false discovery rate-adjusted p; Significance was set at $P \leq 0.05$ ; MDD, major depressive disorder; PTSD, post-traumatic stress disorder; ADHD, attention deficit hyperactivity disorder.

### Gene-based analysis

Gene-based analysis was used to identify genes containing potential association signals that were not highlighted by the individual SNP analysis, but which might nevertheless contribute to biological mechanisms underlying suicidality. Gene-based analysis highlighted *CNTN5, ADCK3/COQ8A*, *CEP57* and *FAM76B* and *DCC* for suicidality (primary analysis, Supplementary Figure 5), *EIF4A1* and *SENP3* and *DCC* for DSH (Supplementary Figure 6) and *CDKAL1, CNTN5,* and *ADCK3/COQ8A* for SIA (Supplementary Figure 7). Regional plots for these genes (except for *CNTN5,* which was also identified in the SNP-based analysis) are presented in Supplementary Figure 8).

### Biology of the suicidality-associated loci

Suggested functions of genes within the suicidality-associated loci are presented in Supplementary Table 5 and the Supplementary Results. Notable findings were the chromosome 11 locus located within *CNTN5*, a very large gene expressed predominantly in brain in adults (Figure 2D); eQTL analysis supporting the gene-based analysis to highlight *CEP57* as another potential candidate gene on chromosome 11 (Supplementary Table 6) as well as several candidate genes on chromosome 9 (Supplementary Figure 9 and Supplementary Results); and gene-based analysis identifying *DCC* as a potential candidate gene. The loci and genes within these loci have been previously associated with a variety of relevant traits (Supplementary Table 7). Of the SNPs with suggestive evidence for association with suicidal behaviour, 21 were available in this study and four demonstrated nominal (p<0·05) association with suicidality in this study (Supplementary Table 8). Only for rs72940689 (14) was the effect direction consistent with the previous report.

## Discussion

### Main findings

Using a very large population-based cohort, we identified multiple genetic loci associated (at genome-wide significance) with suicidality. We found that increased genetic burden for suicidality was associated with increased risk of completed suicide within an independent sub-sample of UK Biobank and we found consistent genetic correlations with a range of psychiatric disorders and psychopathological traits, particularly MDD and anxiety disorders. Separate analyses of DSH and SIA identified one additional signal (for SIA) and suggested that the genetic architecture of DSH is likely to be distinct from that of SIA. Whilst not a true replication, the consistent effect size and direction (and borderline nominal significance) of rs598046-G (*CNTN5*) on suicide attempts in an independent cohort is of interest.

Most previous genetic studies of suicidal behaviour and completed suicide (Supplementary Table 8) been in subjects with known diagnoses of major mental illness. As such, it is difficult in these studies to separate genetic variants for suicidality *per se* from the risk variants for psychiatric disorder. In addition, the limited overlap between suicidality-associated loci identified across these previous studies likely reflects substantial differences in recruitment protocols, participant characteristics (including psychiatric diagnoses) and the variation in the method of assessment of suicidal behaviour. Recently, a general population study in Denmark of suicide attempts identified genetic loci at genome-wide significance (14). Our study extends this approach by investigating a broader phenotype in the primary analysis, as well as the specific impact of DSH versus SIA in secondary analyses. In line with a Research Domain Criteria (RDoC) approach, we used the full spectrum of suicidal thoughts and behaviours assessed within a predominantly non-clinical sample. The fact that increased genetic burden for suicidality was associated with increased risk of completed suicide in a separate sub-sample represents an important validation of our approach to the suicidality phenotype. The genetic correlation with MDD was strong, but the incomplete overlap supports the hypothesis that part of that predisposition to suicidal ideation and behaviour may be distinct from genetic predisposition to depression(25)

In line with our evidence that suicidality has a polygenic component, inclusion of more SNPs (using more relaxed P-thresholds) within a PRS typically demonstrates more significant effects in contrast to strict thresholds because more information and power is provided by including the greater number of SNPs used. Currently there is no agreed threshold that should be considered in these analyses, therefore we reported several PRS analyses. Future work on PRSs for suicidality should seek to identify thresholds that optimally facilitate stratification of clinical and non-clinical populations.

### Biology

Known biology of the suicidality-associated loci (Supplementary Results) highlights three particularly interesting potential candidate genes: *CNTN5*, *CEP57* and *DCC*. *CNTN5* encodes contactin 5 (also known as NB-2), which is a good functional candidate. CNTN5 is a glycosylphosphatidylinositol (GPI)-anchored extracellular cell adhesion protein of the immunoglobulin superfamily, thought to have a role in the formation and maintenance of brain circuitry(26). Centrosomal protein of 57 kDa (CEP57, encoded by *CEP57*) is important for in cell division, with loss of function variants causing a mosaic variegated aneuploidy syndrome, which can include brain abnormalities and mental retardation (OMIM #607951 and #614114). The netrin 1 receptor (encoded by *DCC*) has been robustly associated with depression(27), schizophrenia(28) and related traits. Speculation as to the manner in which variation in these genes act to influence these related traits is hindered by incomplete understanding of the functions of these genes in the brain during development and aging.

### Strengths and Limitations

We acknowledge some limitations to this work. We appreciate that this study is unable to distinguish between putative subtypes of suicide, such as stress-responsive and non-stress-responsive suicidality(29). In addition, we cannot discount the role of study selection or recall bias within the UK Biobank data. Survivor bias might also influence our findings, due to the age at recruitment, however this would likely lead to more conservative effect estimates. We also recognise that the questions used to create the ordinal suicidality phenotype means that some individuals with non-suicidal self-harm behaviours might be included (specifically questions H2 and H3), although this would dilute, rather than inflate, our findings. Our SNP heritability estimates were similar to those reported for other complex psychiatric phenotypes (such as MDD(30)) and the overlap with clinically relevant phenotypes (at the levels of loci, PRS and whole-genome genetic correlations) all suggest that our findings are robust.

### Implications for future work

This study highlights a component of suicidal predisposition that is distinct from MDD predisposition and the potential relevance of *CNTN5, CEP57* and *DCC* to suicidality, further study of which may provide valuable insight into the underlying biology of suicide. Genetic vulnerability to suicide is of course likely to be only a small part of the overall pathophysiology of what is clearly a highly complex and clinically and psychologically heterogeneous phenotype. A major current challenge for the field of suicide research is to integrate new discoveries on the genetics of suicide with known psychiatric, social, psychological and environmental risk factors (such as poverty, substance misuse and childhood sexual abuse), to develop more sophisticated models of risk, and ultimately to develop genetically-informed social, psychological and public health interventions.

### Conclusions

In the largest GWAS to date of suicidality we identified several new candidate genes that may be relevant to the biology of completed suicide. We also demonstrated substantial genetic correlation between suicidality and a range of psychiatric disorders and, by finding an association between genetic loading for suicidality and completed suicide, we provide preliminary evidence for the potential utility of PRSs for patient and population stratification. We hope these discoveries will facilitate new avenues of research on this complex but important phenotype.

## Acknowledgements

We thank all participants in the UK Biobank study. UK Biobank was established by the Wellcome Trust, Medical Research Council, Department of Health, Scottish Government and Northwest Regional Development Agency. UK Biobank has also had funding from the Welsh Assembly Government and the British Heart Foundation. Data collection was funded by UK Biobank. RJS is supported by a UKRI Innovation‐ HDR-UK Fellowship (MR/S003061/1). JW is supported by the JMAS Sim Fellowship for depression research from the Royal College of Physicians of Edinburgh (173558). AF is supported by an MRC Doctoral Training Programme Studentship at the University of Glasgow (MR/K501335/1). KJAJ is supported by an MRC Doctoral Training Programme Studentship at the Universities of Glasgow and Edinburgh. CLN is supported by a UKRI Innovation Fellowship (MR/R024774/1). DJS acknowledges the support of the Brain and Behavior Research Foundation (Independent Investigator Award 1930), a Lister Prize Fellowship (173096) and the MRC Mental Health Data Pathfinder Award (MC_PC_17217).

The funders had no role in the design or analysis of this study, decision to publish, or preparation of the manuscript.

## Author contributions

Study concept and design: Smith, Strawbridge, Ward, Graham, Shaw, Cullen Acquisition, analysis, or interpretation of data: Strawbridge, Ward, L.Lyall, Niedzwiedz, Langan Martin, D.Lyall, Bailey and Smith

Drafting of the manuscript: Strawbridge, Pearsall, Bailey and Smith

Critical revision of the manuscript for important intellectual content: All authors Statistical analysis: Strawbridge, Ward, Graham, Shaw and Ferguson

Administrative, technical, or material support: Pell and Smith

Study supervision: Smith

